# The dominant lineage of *Phakopsora pachyrhizi* in the United States of America does not have a Brazilian origin

**DOI:** 10.1101/2025.01.26.634911

**Authors:** Everton Geraldo Capote Ferreira, Yoshihiro Inoue, Harun M Murithi, Tantawat Nardwattanawong, Jitender Cheema, Ruud Grootens, Sirlaine Albino Paes, George Mahuku, Matthieu H A J Joosten, Glen Hartman, Yuichi Yamaoka, M Catherine Aime, Sérgio H Brommonschenkel, H Peter van Esse, Yogesh K Gupta

**Affiliations:** 2Blades, Evanston, Illinois, USA; The Sainsbury Laboratory, University of East Anglia, Norwich, UK; International Institute of Tropical Agriculture (IITA), Dar es Salaam, Tanzania; Laboratory of Phytopathology, Wageningen University, Wageningen, The Netherlands; Department of Computational and Systems Biology, John Innes Centre, Norwich Research Park, Norwich NR4 7UH, UK; Departamento de Fitopatologia, Universidade Federal de Viçosa, Viçosa, Brazil; Department of Crop Sciences, University of Illinois, Urbana, IL, 61801, USA; Institute of Life and Environmental Sciences, University of Tsukuba, Japan; Department of Botany and Plant Pathology, Purdue University, West Lafayette, IN 47907, USA

**Keywords:** Fungicide Resistance, Lineages, Soybean Rust, Population Structure, Phakopsoraceae, Pucciniales

## Abstract

- Asian soybean rust (ASR), caused by the obligate biotrophic fungus, *Phakopsora pachyrhizi,* was first reported in the continental United States of America (USA) in 2004 and over the years has been of concern to soybean production in the USA. The prevailing hypothesis is that *P. pachyrhizi* spores were introduced into the USA via hurricanes originating from South America, particularly Hurricane Ivan.
- To investigate the genetic diversity and global population structure of *P. pachyrhizi,* we employed exome-capture based sequencing on 84 field isolates collected from different geographic regions worldwide. We compared the gene-encoding regions from all these field isolates and found that four major haplotypes are prevalent worldwide. Here, we provide genetic evidence supporting multiple incursions that have led to the currently established *P. pachyrhizi* population of the USA. Phylogenetic analysis of mitochondrial genes further supports this hypothesis.
- Notably, we observed limited genetic diversity in *P. pachyrhizi* populations in Brazil, suggesting a clonal population structure in that country that contrasts to populations from the USA and Africa.
- This study provides the first comprehensive characterization of *P. pachyrhizi* population structures defined by genetic evidence from populations across major soybean growing regions.

## Introduction

Global food security is threatened by emerging and re-emerging pests and pathogens (Ristaino *et al*., 2021). Recent incursions of plant diseases such as wheat blast in Bangladesh (Islam *et al*., 2016), wheat stem rust in Western Europe (Saunders *et al*., 2019) and Fusarium wilt caused by Tropical Race 4 in Venezuela and Peru (Acuña *et al*., 2022) represent serious threats to crop productivity. Despite the use of modern agricultural practices, 11–30% of crops are still estimated to be lost due to microbial diseases and pests (Savary *et al*., 2019). Soybean is one of the primary sources of edible oil and plant proteins. Asian soybean rust (ASR), caused by the obligate biotrophic fungus *Phakopsora pachyrhizi* Syd. & P. Syd. is a highly destructive disease of soybean (Kelly *et al*., 2015). In Brazil, the cost of managing ASR was estimated at up to US $2.2 billion during the 2013/2014 growing season (Godoy *et al*., 2016). However, in recent years, the economic impact of this disease has declined due to the implementation of effective public policies, such as regulated sowing dares, the adoption of a host-free period, and changes in cropping systems. These measures have helped mitigate environmental conditions favourable for ASR epidemics (Godoy *et al*., 2016).

*P. pachyrhizi* was first reported in Japan in 1902, and until 1934 the disease was only reported across Asia and Australia (Ono *et al*., 1992). In 1994, ASR was first reported in Hawaii (Killgore & Heu, 1994). From 1996–2001, the pathogen was reported in various regions across Southern and Central Africa (Levy, 2005). However, unconfirmed *P. pachyrhizi* occurrences in Africa before 1996 were also reported (Javaid & Ashraf, 1978; Haudenshield & Hartman, 2015). In 2001, *P. pachyrhizi* was reported in Paraguay and Brazil (Rossi, 2003; Yorinori *et al*., 2005) and the pathogen quickly spread across South America over the following three years. However, ASR was not reported in mainland USA until 2004, when it was reported for the first time in Louisiana (Stokstad, 2004; Schneider *et al*., 2005). Shortly after this, the presence of *P. pachyrhizi* was detected across the Southeast USA in Alabama, Arkansas, Florida and Mississippi, and it was perceived as a serious threat to soybean production in those regions (Stokstad, 2004). Aerobiological model simulations implicated hurricanes, especially hurricane Ivan, as the most likely mode of introduction of *P. pachyrhizi* urediniospores, originating from South America, into the continental USA in 2004 (Isard *et al*., 2005, 2007; Pan *et al*., 2006). Unfavourable disease conditions (such as the harsh winter climate) along with meticulous monitoring were critical in limiting the impact of the disease (Sikora, 2014; Kelly *et al*., 2015).

Previous studies reporting the genetic diversity of *P. pachyrhizi* populations worldwide have shown a lack of genetic differentiation and population structure between populations from South America, particularly Brazil and Argentina (Jorge *et al*., 2015; Darben *et al*., 2020; Rocha *et al*., 2024) and Nigeria (Twizeyimana *et al*., 2011). On the contrary, this limited genetic diversity does not necessarily correlate with a low diversity of virulence profile within these populations of *P. pachyrhizi*. In fact, several unique and shared pathotypes have been identified across different countries including Brazil, Argentina and Paraguay (Akamatsu *et al*., 2013, 2017), the USA (Walker *et al*., 2011; Paul *et al*., 2015), Kenya, Malawi, and Nigeria (Twizeyimana *et al*., 2009; Murithi *et al*., 2017, 2021), Uruguay (Stewart *et al*., 2019; Larzábal *et al*., 2022), Mexico (García-Rodríguez *et al*., 2022), and Bangladesh (Hossain & Yamanaka, 2019). These virulence profile studies suggest high levels of gene flow between *P. pachyrhizi* populations across large geographic regions and continents, indicating the long-distance dispersal of *P. pachyrhizi* spores. Interestingly, studies of early *P. pachyrhizi* isolates collected from the 2000’s in the USA (Zhang *et al*., 2012) and Brazil (Freire *et al*., 2008) revealed a high genetic diversity during the initial outbreaks of ASR. These two studies suggest multiple ribotypes (a pattern of ribosomal RNA bands, to detect polymorphism) in Brazilian populations, originating from Africa and Asia, while USA populations contained ribotypes from South America, Africa, Asia, and Australia. These findings suggest that early *P. pachyrhizi* populations in both Brazil and the USA were established through multiple introductions. Although all previous studies had used molecular markers (SSRs and AFLP markers), as well as housekeeping gene sequences (*ITS* and *ADP* genes), these approaches have their own advantages and limitations (Rush *et al*., 2019; Sheeja *et al*., 2021). More importantly, although these housekeeping genes are not under strong selection pressure, they do not provide a broad picture of the genetic differences at the whole genome level. Therefore, inferences on the global migration of *P. pachyrhizi* based on these analyses might be incomplete.

The 1.25 Gb genomes of three *P. pachyrhizi* isolates were recently sequenced (Gupta *et al*., 2023), facilitated by advances in long-read sequencing technologies. The *P. pachyrhizi* genome, characterized by its large size, high repeat content (93% transposable elements), significant heterozygosity, and the dikaryotic nature of infectious urediospores, has historically presented significant challenges for genome assembly and robust population genomic analyses. The availability of these high-quality genome assemblies now enables population studies *P. pachyrhizi* with much greater resolution, comparable to what has been applied to other plant pathogens. New approaches such as field-pathogenomics has emerged as a powerful approach for pathogen diagnostics, surveillance, evolutionary analysis, and analysis of population structure of plant pathogens, as has been performed for wheat stripe rust, wheat powdery mildew, wheat blast and *Fusarium* head blight (Hubbard *et al*., 2015; Kelly & Ward, 2018; Jouet *et al*., 2019; Thilliez *et al*., 2019; Radhakrishnan *et al*., 2019).

In this study, we applied phylogenetic and population genetic approaches to investigate the global spread of *P. pachyrhizi*, and its implications on the population structure and genetic diversity of *P. pachyrhizi* populations in the two major soybean producing countries. We sequenced 84 field isolates collected from soybean-growing regions of East Africa, South America, North America, Asia, and Australia. We found mutations in mitochondrial genes, suggesting that the soybean rust population in the USA, Japan and Australia potentially originated from a common ancestor. This hypothesis is further supported by polymorphisms identified in nuclear gene sequences. Notably, both the Brazilian and African *P. pachyrhizi* populations have low genetic diversity when compared to the USA population, suggesting a strong bottleneck for selection in Brazil and Africa. Our data furthermore indicated high levels of genetic diversity in USA populations, with clustering patterns suggesting multiple incursions of *P. pachyrhizi* into the continental USA. Our results demonstrate that the current view on the spread of *P. pachyrhizi* is still incomplete but does not support a sole South American hypothesis for the origin of ASR in the USA, but rather, highlight a complex migration dynamic of *P. pachyrhizi* populations.

## Results

To estimate the genetic diversity of *P. pachyrhizi,* infected soybean samples were collected from different geographic regions (**Fig. 1; Table S1**). A bait library was designed to capture *P. pachyrhizi* gene sequences from total DNA extracted from the infected samples based on RNA-seq data from a time course study of the UFV02 isolate of *P. pachyrhizi* (Gupta *et al*., 2023). In total, 84 field isolates were sequenced, and an uninfected soybean cv. Williams 82 (W82) leaf sample was included to assess the specificity of the baits. All field isolates were collected from infected soybeans, except four field isolates from the USA and the Japanese K1-2 isolate (Yamaoka *et al*., 2014), which were collected from kudzu (*Pueraria lobata* (Willd.) Ohwi), a wild legume host of *P. pachyrhizi*.

**Figure 1.**
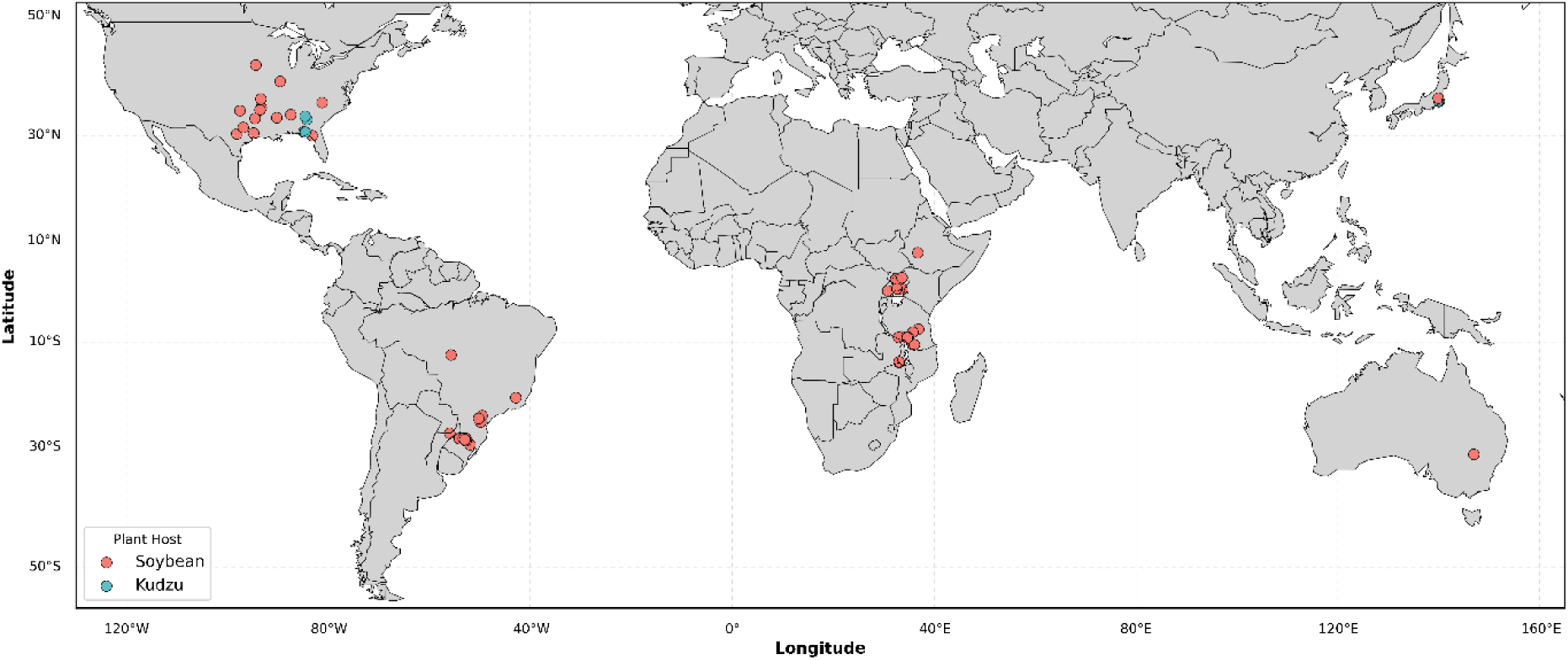
Sampling locations of *P. pachyrhizi* analysed by exome-capture sequencing. Dots on the map represent the locations and numbers of the samples collected. The color corresponds to the host where the samples/isolates originated from.

To evaluate the performance of the bait library, we mapped the reads of UFV02 to its reference genome assembly (Gupta *et al*., 2023). The UFV02 genome contains 22,467 annotated genes, of which 10,942 were captured (**Fig. 2a**). This outcome was anticipated, as the bait library was specifically designed to target only expressed genes. Indeed, we captured 7,822 out of 9,437 expressed genes described in the UFV02 transcriptome, suggesting that 82.9% of the expressed genes from UFV02 were successfully captured using the exome-capture based method (**Fig. 2b**). We then assessed the sequence read coverage for the 10,942 captured genes across all 84 field samples, and from the uninfected leaf of soybean W82. As expected, no *P. pachyrhizi* reads were detected in the W82 control, demonstrating the high specificity of the exome-capture method for *P. pachyrhizi* DNA sequences. We observed differences in read-depth and coverage in the nuclear genome, as well in the mitochondrial genome across all the samples, which is probably related to the severity level of *P. pachyrhizi* infection (**Fig. S1 & Fig. S2**).

**Figure 2.**
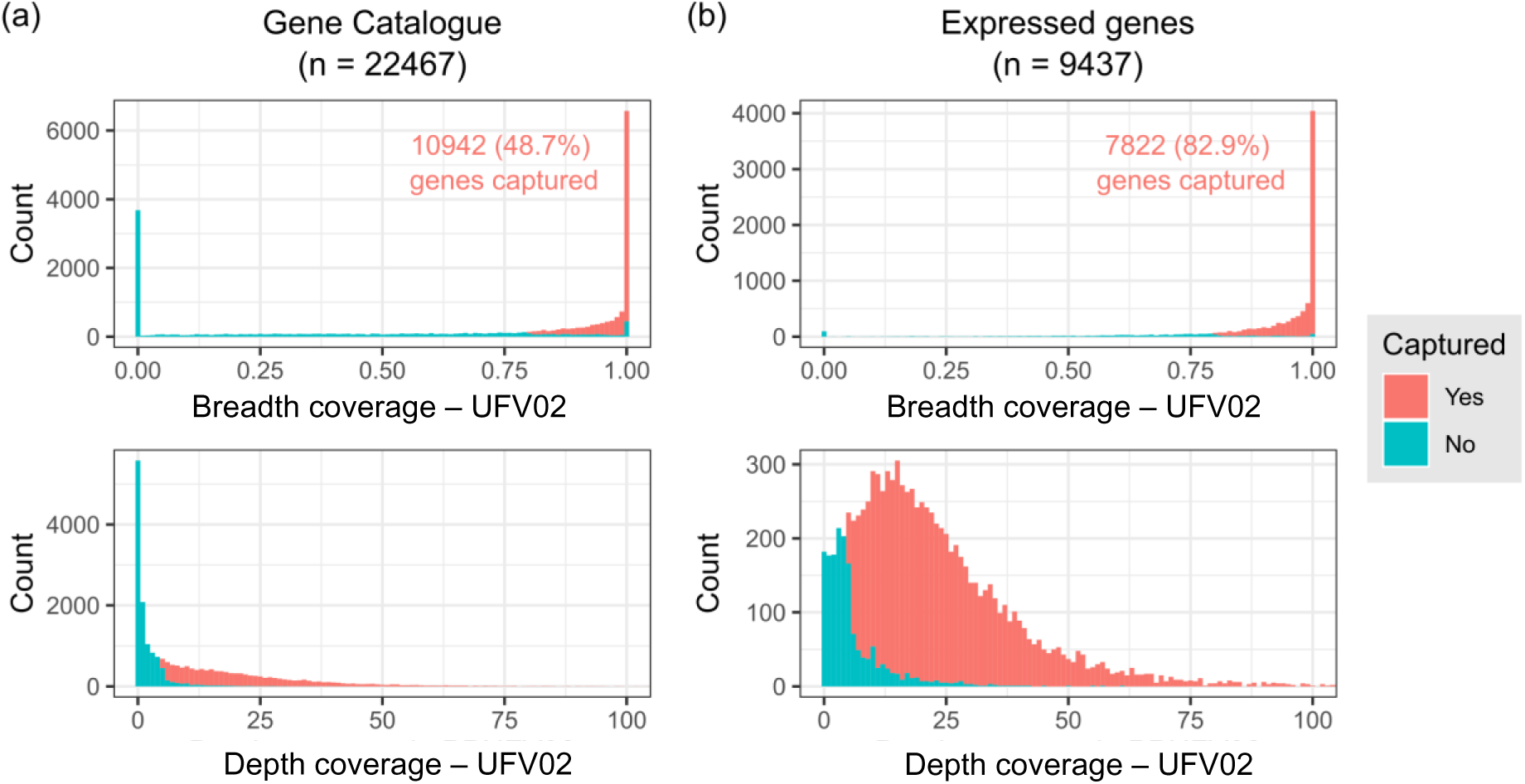
**Evaluation of the gene space coverage by exome-capture sequencing of *P. pachyrhizi*.** Histograms representing the breadth (upper panel) and depth (lower panel) coverage of *P. pachyrhizi* genes by exome-capture sequencing reads of a *P. pachyrhizi* isolate UFV02. The whole gene set (UFV02 “Gene Catalogue”) and a set of expressed genes were evaluated in (**a**) and (**b**), respectively. A gene was judged “captured” if the gene had a breadth coverage of over 0.8 and a depth coverage of over 5.

### Distinct mitochondrial haplotypes reveal the population structure in *P. pachyrhizi* at the geographic level

Mitochondrial (mt) genomes evolve independently from nuclear genomes, and polymorphisms in introns and intergenic regions of conserved mitochondrial genes are often used to assess the genetic diversity within populations (Ballard & Whitlock, 2004; Pantou *et al*., 2008; Kim *et al*., 2016; Jiménez-Becerril *et al*., 2018). To evaluate the polymorphisms in the mt genomes of the field isolates, we mapped their reads to the mitochondrial genome of *P. pachyrhizi* (Stone *et al*., 2010).

Although the *P. pachyrhizi* mt genome is highly conserved across all field isolates, we identified six SNPs and four InDels between the isolates. Based on those variants, four major haplotypes were observed across the 84 sequenced field isolates (**Fig. 3a**). Notably, a non-synonymous SNP marker (a T to C change at position 15,470) resulting in an F129L mutation in the cytochrome b (CYTB) gene was identified. This mutation, predominantly found in field isolates from South America, is exclusive to haplotype II (**Fig. 3a**). This F129L mutation, previously reported in South America, has been linked to reduced sensitivity to Quinone outside inhibitor fungicides (QoIs) (Klosowski *et al*., 2016b; Müller *et al*., 2021). Remarkably, this mutation was also observed in two field isolates from Uganda rendering this the first report of the F129L mutation in *P. pachyrhizi* outside South America (**Fig 3a**).

**Figure 3.**
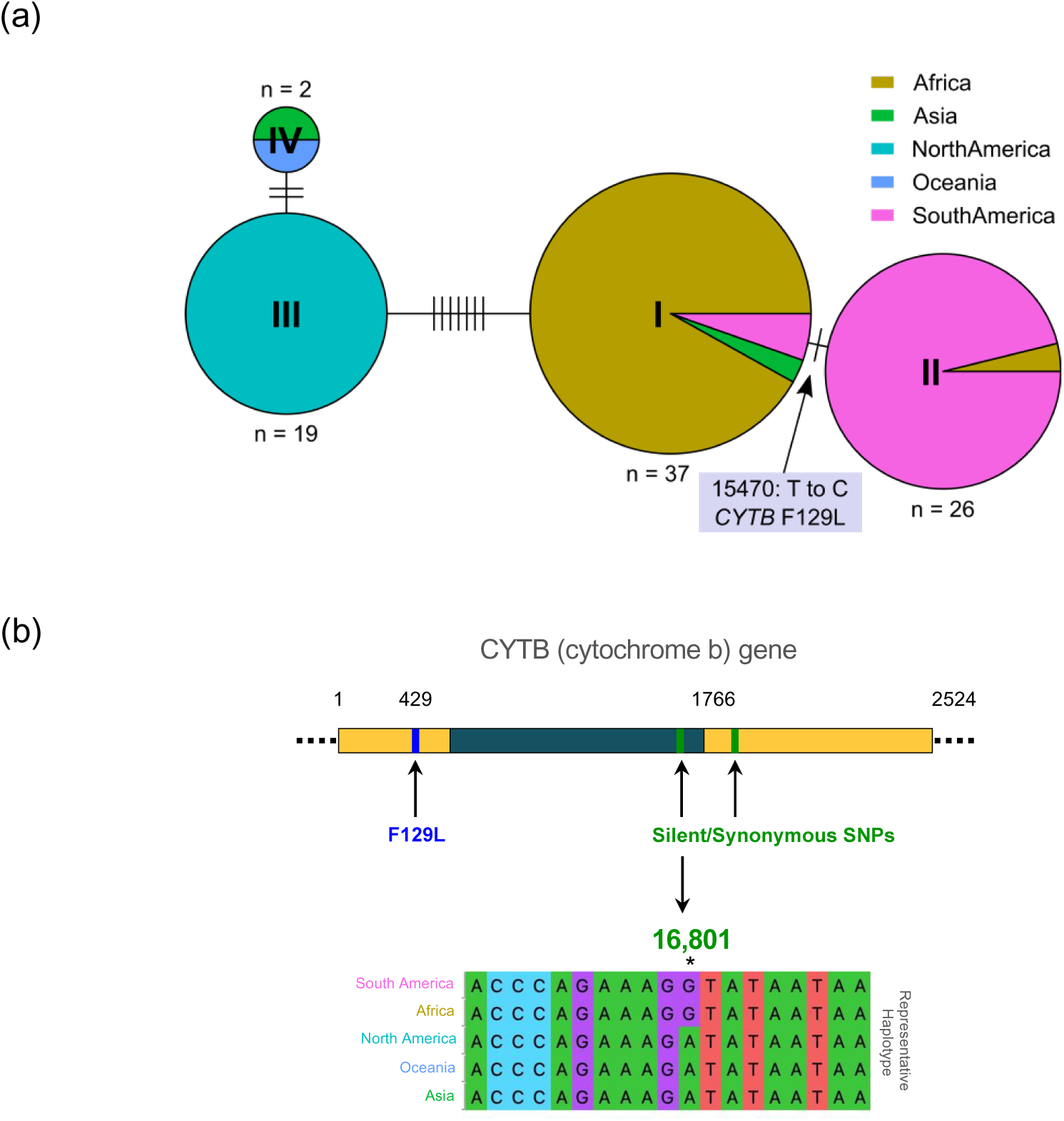
Haplotype network inferred from nucleotide polymorphisms in the mitochondrial genome. **(a)** Pie charts represent haplotypes and their geographic origins. The size of each node is proportional to the number of isolates sharing that haplotype. Connecting edges indicate genetic relationships, with each tick mark representing a single SNP or INDEL. (**b**) A non-synonymous SNP (F129L) associated with resistance to quinone outside inhibitor (QoI) fungicides is marked by a blue vertical bar. Synonymous SNPs (no amino acid change) are marked by green vertical bars. Representative haplotypes surrounding SNP marker 16,801 (converted to KASP assay), which differentiate populations based on geographic location, are highlighted in green.

Surprisingly, all field isolates from the USA clustered into a single haplotype, haplotype III, exhibiting no intragenic variation. The USA field isolates showed seven polymorphic sites (alternative alleles) compared to those from South America and East Africa (**Fig. 3a**). We identified two unique mutations (synonymous SNPs) in the CYTB gene, present in all the USA field isolates and in the K1-2 isolate from Japan, distinguishing haplotype III from haplotypes I (mainly African field isolates) and II (mostly South America) (**Fig. 3b**). This suggests that the current USA population emerged from a single, distinct lineage separate from the South American and East African *P. pachyrhizi* populations. The low genetic variation observed in the mt genome indicates limited genetic exchange between populations, as expected, considering that *P. pachyrhizi* spreads through clonally produced urediniospores. Lastly, the mitochondrial haplotype structure reveals clear geographic separation of *P. pachyrhizi* populations.

### Unique origin of the *P. pachyrhizi* lineage in the USA

To elucidate the population structure of *P. pachyrhizi*, we mapped the sequencing reads from all the field isolates to the UFV02 reference genome assembly. Based on read depth coverage, we selected 53 field isolates and identified 33,634 high-confidence SNP markers to define the population structure and identify genetically related *P. pachyrhizi* isolates. We conducted a discriminant analysis of principal components (DAPC), which revealed four distinct *P. pachyrhizi* populations, each corresponding to a different continent. An unsupervised genotype clustering analysis further corroborated the presence of these four well-supported clusters (**Fig. 4a & b**).

**Figure 4.**
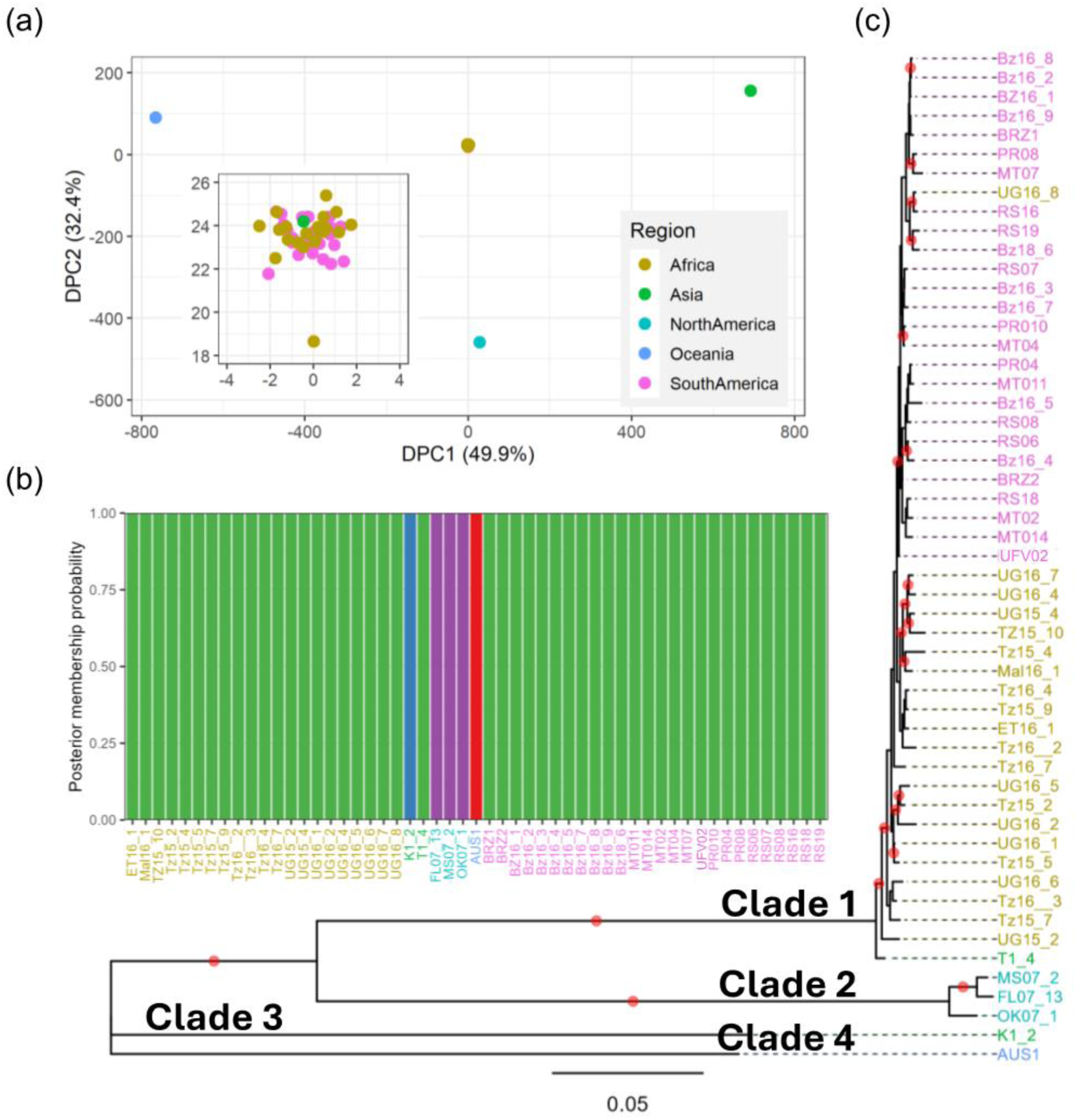
Population structure and phylogenetic relationships of *P. pachyrhizi* samples inferred from nuclear genome SNPs. (**a**) Discriminant Analysis of Principal Components (DAPC) using SNP markers from 53 samples, with principal components 1 and 2 plotted. (**b**) Probability of population membership is shown on the y-axis for each sample, grouped in bins along the x-axis. The stacked bar chart represents assigned population membership probabilities, with clusters distinguished by different colours. (**c**) Phylogenetic tree inferred from 33,634 SNP markers. Nodes with bootstrap support greater than 80% are marked with red dots.

To assess evolutionary relationships among field isolates and to determine the extent of clonality, we constructed a maximum-likelihood (ML) phylogenetic tree using the 33,634 biallelic SNP markers. The phylogenetic analysis grouped the *P. pachyrhizi* field isolates into four major clades (**Fig. 4c**), consistent with the clusters identified in the DAPC analysis (**Fig 4a**). The largest clade, Clade 1, includes field isolates from both South America and East Africa. Within Clade 1, these isolates further segregated into two sub-groups according to their geographic origin, suggesting a lack of genetic exchange between *P. pachyrhizi* populations in South America and East Africa, and implying that these populations may be evolving independently.

All USA field isolates formed a distinct clade, Clade 2, separate from Clade 1, with strong bootstrap support (>80%) (**Fig. 4c**). Given the high degree of heterozygosity observed in the *P. pachyrhizi* genome, we assessed heterozygosity across all field isolates (**Fig. S3**). Notably, USA field isolates, the Oceanic isolate, AUS1, and the Japanese kudzu isolate, K1-2, exhibited higher heterozygosity (17.2– 28.0% of the SNP sites showed a heterozygous genotype) compared to other field isolates (6.4–12.6% of the SNP sites showed heterozygous genotype; **Fig S3**). These findings, combined with haplotype analysis, suggest that the *P. pachyrhizi* population in the USA is not genetically linked to the South American population. Additionally, our phylogenetic analysis revealed that the K1-2 isolate (collected from the kudzu host in Japan) and the AUS1 isolate (collected from soybean in Australia), diverged significantly from all the other sequenced isolates, forming separate clades (clade 3 and clade 4, respectively). These results suggest that *P. pachyrhizi* populations in Japan and Oceanic regions are highly diverse and warrant further investigations.

For this study, we used USA field isolates collected in 2007–2008, and two samples from Japan (T1-4 and K1-2). To investigate the genetic relationship with the current *P. pachyrhizi* population in the USA (2017) and in Japan (2012–2017), we converted the two SNP makers from the CYTB gene that efficiently separated the samples based on the geographic origin (**Fig. 3**) into competitive allele-specific PCR (KASP) assays (**Table S2)**. For this approach, one of the KASP assays (COB2_2, 16,801) showed clear allelic discrimination and was kept for future analyses. The KASP results showed that the recent USA *P. pachyrhizi* population follows a similar pattern (same allele detected) to the earlier isolates from 2007 and are not genetically linked to the South American population (**Table S3**). In summary, the KASP assay COB2_2 provides a valuable resource for monitoring pathogen dispersal on a global scale, as well as helps to define the genetic structure of the pathogen population.

## Discussion

Current population studies of *P. pachyrhizi* have primarily relied on simple sequence repeat (SSR) markers, amplified fragment length polymorphisms (AFLPs), and single-locus sequencing of genes, such as internal transcribed spacer (ITS) and ADP-ribosylation factor (ARF) (Anderson *et al*., 2008, Zhang *et al.,* 2012 and Twizeyimana *et al*., 2011). While these methods have provided valuable insights, they lack the resolution needed to capture fine-scale population structure and genetic diversity. Our findings corroborate previous studies, confirming low levels of genetic diversity within Brazilian and African populations, with comparatively higher diversity in the US *P. pachyrhizi* populations. Although these neutral markers provide valuable baseline data on population differentiation, they offer limited insights into the adaptive genetic structure needed for understanding evolutionary dynamics and pathogenic variability at the genome level. In contrast, whole-genome or reduced representation sequencing approaches offer a higher resolution strategy for studying adaptive evolution and population diversity, especially in understudied pathogens (Hubbard *et al*., 2015; Jouet *et al*., 2019; Thilliez *et al*., 2019).

Like other rust species, *P. pachyrhizi* is an obligate biotrophic pathogen, dependent on living host tissue for growth and propagation. This inability to grow *in vitro* presents significant challenges and limitations for genetic studies and other investigations (Panstruga, 2003; Lorrain *et al*., 2019). Additionally, the large genome size of *P. pachyrhizi* (1.25 Gb), and its high repeat content (93% of TEs) further complicate large-scale population studies (Gupta *et al*., 2023). These challenges stem from the substantial resources required to isolate high-molecular-weight DNA, the high costs of sequencing, and the complexity of analyzing such repetitive and heterogeneous genomic data. To overcome these limitations, we developed an exome-capture-based approach that selectively targets genic regions. This approach have been extensively used for genetic studies in plants (King *et al*., 2015; Mukrimin *et al*., 2018; Hussain *et al*., 2018; Dong *et al*., 2020), human pathogens (Carpi *et al*., 2015; Quek & Ng, 2024), animals (McClure *et al*., 2014; Fairfield *et al*., 2015) and plant pathogens such as *Phytophthora infestans* (Thilliez *et al*., 2019; Coomber *et al*., 2024). Using the recently sequenced *P. pachyrhizi* genome as a scaffold, we validated the specificity and efficacy of this approach, successfully capturing 83% of all expressed genes in a proof-of-concept study. This significantly enhanced our ability to draw accurate inferences on pathogen migration and population structure of *P. pachyrhizi*.

### Distinct mitochondrial haplotypes and global lineages of *P. pachyrhizi*

Through mt genome analysis, we identified four distinct haplotypes among *P. pachyrhizi* field isolates. A single non-synonymous mutation, F129L, in the CYTB gene distinguished haplotypes I and II, with haplotype II carrying the F129L mutation in 95% of the South American field isolates. The reduced efficiency of various fungicide classes in controlling ASR in Brazil has been associated with the increasing prevalence of known mutations in fungicide target genes (Godoy, 2012; Godoy *et al*., 2022). In particular, the F129L mutation is associated with reduced sensitivity to QoI fungicides (Godoy *et al*., 2016), and has been previously observed at high frequencies in South American *P. pachyrhizi* populations (Klosowski *et al*., 2016b,a; Müller *et al*., 2021; Claus *et al*., 2022). Notably, we detected the F129L mutation in two field isolates from Uganda, making the first report of this mutation outside South America. Although strobilurin-based fungicides remain effective against ASR in African countries such as Uganda and Ethiopia (Kawuki *et al*., 2003; Abebe *et al*., 2022), the detection of this mutation underscores the need for continuous monitoring.

Interestingly, USA field isolates formed a unique haplotype, haplotype III, with seven unique polymorphic sites compared to haplotypes I and II. The *P. pachyrhizi* field isolates from the USA analyzed in this study did not exhibit the F129L mutation. This absence is likely due to the relatively low ASR incidence in the USA, which reduces the need for fungicide applications and, consequently, the selective pressure for this mutation (Bradley *et al*., 2021). This genetic uniqueness suggests limited selection pressure for QoI resistance, as ASR outbreaks in the USA are sporadic and often constrained by climatic factors. This represents the first report identifying a unique genetic signature in USA *P. pachyrhizi* populations, suggesting a distinct evolutionary trajectory.

The emergence of new races and pathotypes in plant pathogenic fungi often depends on their mutation and recombination rates (Wyka *et al*., 2022; Amezrou *et al*., 2024). In *P. pachyrhizi*, long-distance dispersal via airstreams can promote genetic exchange, particularly when spores land on an alternate host. This process can significantly influence population substructure, even in geographically separated regions. Our phylogenetic reconstruction revealed two major clades: Clade 1, comprising isolates from South America and Africa, and Clade 2, exclusively containing USA isolates. The clustering of South American and African field isolates into Clade 1 supports the previous hypothesis that Brazilian *P. pachyrhizi* populations likely originated from Africa (Freire *et al*., 2008). Our results showed that, despite its airborne nature, the genetic similarity between populations from these two regions suggests a predominantly clonal structure, likely facilitated by similar tropical climates and asexual reproduction (Goellner *et al*., 2010).

In contrast, the USA population, represented by Clade 2, exhibited significant genetic divergence, potentially driven by the overwintering of *P. pachyrhizi* on alternative plant hosts such as kudzu in regions like Florida and the Gulf Coast, where milder conditions prevail (https://soybean.ipmpipe.org/soybeanrust/). During harsh winters in the USA Midwest, the absence of soybean as a host is mitigated by the presence of kudzu, which provides refuge for this obligate pathogen. However, the harsh winter conditions likely impose a bottleneck effect, reducing the survival of *P. pachyrhizi* spores and further shaping its population structure(Harmon *et al*., 2005; Isard *et al*., 2005, 2007; Kelly *et al*., 2015). Each year, airborne spores are carried by wind from southern regions to the Midwest, but their late arrival in the growing season reduces the likelihood of severe ASR outbreaks. This annual spore movement, combined with the pathogen’s dependence on kudzu as a winter reservoir, likely contributes to the genetic differentiation of USA *P. pachyrhizi* populations from those in South America and Africa.

Additionally, multiple independent introductions of *P. pachyrhizi* into the continental USA may have further shaped the genetic structure of these populations. It is plausible that *P. pachyrhizi* populations of Asian origin entered the USA, potentially via Hawaii and subsequently adapted to infect soybean and kudzu, while surviving the harsh winter conditions. Notably, Japan experiences severe winters, which may explain the shared genetic adaptations observed between Japanese and USA isolates (Twizeyimana & Hartman, 2012; Yamaoka *et al*., 2014). Our results further support this hypothesis as the Japanese and USA isolates from kudzu shared the same mutation in the CYTB mitochondrial gene.

A recent study by da Rocha *et al*., (2024) also identified two major clades within *P. pachyrhizi* populations, corresponding to older and more recent lineages. However, their study was limited by the inclusion of only two USA isolates: one from Hawaii (HW94-1, isolated in 1994) and one from the continental USA (FL07-01, isolated in 2007). Interestingly, FL07-01, clustered closely with isolates from Africa and Brazil, while HW94-1, clustered with older isolates from India, Taiwan and Australia (da Rocha *et al*., 2024). The limited sampling may have underestimated the genetic diversity and population structure within the USA. However, it is plausible that during the initial years of *P. pachyrhizi*’s colonization in the continental USA, multiple introductions occurred with FL07-01 potentially representing a South American lineage that later diminished as populations introduced via Hawaii became dominant.

Our findings challenge the long-held assumption that USA ASR outbreaks stemmed solely from hurricane-driven spore dispersal from Brazil. Instead, the genetic uniqueness and tight clustering of the USA *P. pachyrhizi* population in Clade 2 suggest that *P. pachyrhizi* introductions into the USA may have occurred through independent migration events, rather than a single introduction from Brazil, as previously proposed by Zhang *et al*., (2012).

In addition, isolates from Australia (AUS1) and Japan (K1-2) formed highly divergent clades compared to clades 1 and 2, suggesting substantial genetic differentiation in these populations. The high genetic diversity observed in the USA population is unlikely to arise from somatic hybridization or sexual reproduction, as no alternate host has been identified to date. It is mostly likely that kudzu, a widely distributed alternative host, along with the harsh winters, has driven adaptation in the USA population, enabling it to effectively infect both soybean and kudzu. Interestingly, despite the clonal nature of the *P. pachyrhizi* populations highlighted by the low diversity across different countries, a high diversity in virulence profiles has been consistently reported. Therefore, this indicates genetic forces acting in the selection and generation of different races of the pathogen. Additionally, the role of TEs in fungal genome evolution cannot be overlooked. TEs are known to contribute to adaptive variations in traits such as pathogenicity and virulence (Torres *et al*., 2021; Fouché *et al*., 2022; Oggenfuss & Croll, 2023). With approximately 93% of the *P. pachyrhizi* genome consisting of TEs, of which ∼12% being expressed (Gupta *et al*., 2023), it is plausible that TEs play a critical role in generating genetic variability in the absence of sexual reproduction and somatic hybridization. Recent studies have pointed out the role of TEs in shaping genetic variability in various fungal species. For instance, TEs mediate metal resistance in *Paecilomyces variotii* (Urquhart *et al*., 2022); drive strain-specific evolution and host-specific expression regulation of TEs in *Rhizophagus irregularis* (Oliveira & Corradi, 2024); and contribute to lineage-specific differences in *Magnaporthe oryzae* infecting various grasses (Nakamoto *et al*., 2023). While somatic hybridization has been documented in other rust fungi (Li *et al*., 2019; Sperschneider *et al*., 2023), a mechanism that generates genetic diversity through recombination between nuclei, but has not yet been confirmed in *P. pachyrhizi*. However, a study reported hyphal anastomosis between urediniospore germ tubes in *P. pachyrhizi*, facilitating nuclear migration into a shared hyphal network (Vittal *et al*., 2012). This observation raises the possibility of nuclear exchange and recombination. Although somatic hybridization does not generate new mutations, it can create novel allele combinations from the fusion of two nuclei, potentially driving the rapid emergence of new *P. pachyrhizi* races.

### Implications for Disease Management

Our exome-capture-based sequencing approach provided an in-depth view of genetic variation within the genic regions of *P. pachyrhizi*, enabling the identification of SNP markers associated with the distinct geographic origins. Such genetic markers can readily be converted into low-cost KASP assays, for use in-field monitoring and early detection of fungicide-resistant or more virulent *P. pachyrhizi* populations. Such methodologies will be crucial for enabling on-time targeted interventions, as is being carried out in the case of other important plant pathogens (Bueno-Sancho *et al*., 2017; Radhakrishnan *et al*., 2019; Paineau *et al*., 2024). These methods have great potential for future applications in the detection of resistance-breaking isolates and in monitoring the durability of disease resistance in the field.

## Conclusion

Overall, this study contributes to the understanding of *P. pachyrhizi* population dynamics worldwide and highlights the emergence of a genetically divergent lineage among isolates from the USA, which is likely driven through multiple independent introductions into the country. Exome-capture sequencing technology offers a valuable tool for studying the adaptive genetic variation of this pathogen. Future research should delve into the role of TE’s in generating diversity in the absence of sexual reproduction or alternate host. Additionally, the knowledge of host plant interaction such as kudzu and soybean, under varying climate conditions will be necessary for a more comprehensive understanding of the evolutionary dynamics of *P. pachyrhizi*. These will be important in devising management strategies that can help reduce the continued threat of this highly adaptive and destructive pathogen.

## Materials and Methods

### Field isolates collection

81 ASR infected field isolates from different geographic regions including Australia (n= 1), Argentina (1), Brazil (25), East-Africa (35), and USA (19) (**Table S1**) were collected. The field isolates from East-Africa were collected from Ethiopia, Malawi, Uganda, and Tanzania between 2015 and 2016. The isolates from Brazil were collected in 2015 and 2016 and USA isolates collected in 2007 and 2008. The monopustule isolates UFV02 (Brazil – 2006) (Gupta *et al*., 2023) and T1-4 (Japan – 2007) obtained from cultivated soybean and the monopustule isolate K1-2 obtained from Kudzu (*Pueraria lobata* (Willd.) Ohwi) were also included in the analysis (Yamaoka *et al*., 2014). Spore suspension and inoculation assay were carried out as previously described to obtain infected leaves material for the three monopustule isolates. Briefly, spores of the isolates were heat-shocked at 40°C for 5 min, suspended in an aqueous solution (0.01% Tween 20), mixed thoroughly and concentration adjusted to approximately 5 x 10^4^ spores/mL with a hemocytometer. Four-week-old soybean plants (cv. Williams 82) were sprayed on abaxial surface of the leaves with the spore suspension, kept at 100% relative humidity in the dark for 24 hours. After 24 h, inoculated plants were kept at 22°C and 70% relative humidity, with 16 h photoperiod. 14 days post inoculation (dai), infected leaves were collected and stored at −80°C until DNA extraction. The infection was performed in three biological replicates and for each replicate, three leaves per plant (trifoliates) and three plants were used for each individual replicate.

### Exome capture design, library preparation and sequencing

Genomic DNA from all 84 infected leaf samples (100 mg of leaf tissue) was extracted using the DNeasy Plant Mini Kit (Qiagen Manchester, UK) according to the manufacturer’s instructions. The DNA samples were provided to Arbor Biosciences (Ann Arbor, Michigan, US) for library preparation, target capture enrichment, and Illumina sequencing. Briefly, the target capture baits library was built based on the *P. pachyrhizi* UFV02 transcriptome dataset publicly available (Gupta *et al*., 2023). The probe (also referred here as baits) sequence candidates were designed following the resistance gene enrichment sequencing (RenSeq) pipeline previously established (Jupe *et al*., 2013). The probes sequences were designed over and between exon-exon boundaries of the retrieved expressed gene sequences. The first probe started at the left most nucleotide and followed the predicted coding direction of the gene. The target probe sequences were 100 bp length, 4x tilling and 25 bp overlap between baits. In total around 162,000 baits were designed based on %GC, secondary propensities, cross-complementarity and 80% identity between baits and further synthesized by Arbor Biosciences (Ann Arbor, MI, USA). Genomic DNA from the 84 samples were subjected to target capture hybridization using the biotinylated bait-library. Enriched libraries were submitted for sequencing on an Illumina HiSeq 4000 platform, 150 bp paired-end reads. As a positive control, cv. Williams 82 infected with UFV02 was considered, and as a negative control, DNA of non-inoculated cv. Williams 82 was subjected for the hybridization and sequencing to confirm the cross-hybridization of baits.

### Filtering steps and variant calling

The quality of the raw reads from sequencing was checked using FastQC program (https://www.bioinformatics.babraham.ac.uk/projects/fastqc/). Low quality reads and bases reads were trimmed using the Trimmomatic v0.39(Bolger *et al*., 2014). The filtered paired-end reads were mapped to the *P. pachyrhizi* (UFV02 v2.1) reference genome downloaded from the Joint Genome Institute (JGI) (https://mycocosm.jgi.doe.gov/PpacPPUFV02/PpacPPUFV02.info.html) using BWA v0.7.17 (Li & Durbin, 2009) with the BWA-MEM algorithm and default parameters. Resulting SAM files were sorted and converted into BAM files using SAMtools v1.9 (Danecek *et al*., 2021). The filtered paired-end reads were also mapped to the *P. pachyrhizi* mitochondrial genome (Taiwan 72-1 isolate, GenBank: GQ332420), converted to BAM format and sorted as mentioned above. BAM files generated using nuclear and mitochondrial genomes were used independently to generate variants files for follow-up phylogenetic and population analyses.

For variant calling, PCR duplicates were marked in the BAM files using Picard v1.118 toolkit (http://broadinstitute.github.io/picard/) and variant calling was performed using the GATK in two rounds (Auwera & O’Connor, 2020). First, variants (SNPs and InDels) from all the samples were predicted by HaploTypeCaller function, the resulting VCF was split into two files using SelectVariants function, one for SNPs and another for InDels. Then, VariantFiltration function was applied for hard filtering SNPs with quality thresholds (QD < 2.0; FS > 60.0; MQ < 40.0; MQRankSum < −12.5; ReadPosRankSum < −8.0) and for filtering InDels (QD < 2.0; FS > 200.0; ReadPosRankSum < −20.0; SOR > 10.0). Base quality scores in the BAM files were recalibrated using the filtered variants files using BaseRecalibrator and ApplyBQSR functions. Finally, the recalibrated BAM files were used for the second variant calling using HaplotypeCaller and GenotypeGVCFs functions.

### Phylogenetics and population structure analyses

Only biallelic SNPs from the nuclear genome (missing data < 70%) were used to the phylogenetic analyses. The neighbor-joining trees were inferred using the package phangorn 2.5.5 on R environment using the dist.hamming function with 1,000 bootstrap replicates. The trees were visualized by the online tool iTOL v6 (https://itol.embl.de). The population structure was based on DPAC (Discriminant Analysis of Principal Components), BIC (Bayesian Information Criterion) and Structure analyses. Multivariate analysis using DAPC was performed using the Adegenet package on R environment and the number of genetic clusters showing the optimal BIC number was identified. Estimations of admixture and number of sub-populations were inferred using the Structure software (Pritchard *et al*., 2000). The algorithm ran with a burn-in length of 50,000 and with a simulation length of 100,000 Markov Chain Monte Carlo (MCMC) repetitions. The number of possible genetic cluster (*K*) ranged from 1 to 10, with 20 repetitions per *K*. Variants from the mitochondrial genome were used to build a haplotype network using the HaploNet function in the *pegas* package in R environment. The haplotypes were colored based on the geographic origin and visualized by *ggplot2* package in R.

Breadth and depth of coverage were calculated for all the genes captured by the library. Briefly, breadth of coverage corresponds to the percentage of target bases with at least one mapped read (1x coverage). Proportion of heterozygous/homozygous SNPs for each sample was calculated by bcftools v1.9 (Danecek *et al*., 2021). Nucleotide diversity (π) and Nei’s fixation index (F_st_) between the subpopulations were calculated with vcftools v0.1.17 (Danecek *et al*., 2011) using the whole set of SNPs as well in predicted effectors (Gupta *et al*., 2023) and BUSCO (Simão *et al*., 2015)housekeeping genes (1,335 Basidiomycota single-copy orthologs). Only biallelic SNPs, with a depth coverage (DP) ≥ 3 and max-missing ≤ 0.1 were considered in these analyses.

### Haplotype analysis and KASP validation

In order to validate the mitochondrial SNP markers and uncover further *P. pachyrhizi* genes able to distinguish between ASR subpopulations based on their geographic origins, we have done haplotype analyses using the high-quality SNPs. To perform that, we identified manually those SNPs, checked the gene models they were located, predicted their impact, and checked the expression levels based on public dataset. We developed KASP markers for the selected SNP markers using an in-house python script and we genotyped an extra set of ASR field isolates collected in 2017 from Louisiana, US. DNA was extracted from approximately 100 mg of *P. pachyrhizi s*oybean infected leaves with DNeasy Plant Mini kit (Qiagen, Manchester, UK) following the manufacturer’s protocol. DNA samples were diluted to the concentration of 10 ng/ul and submitted to KASP reaction using PACE^®^ allele-specific master mix (3CR Bioscience, Essex, UK) following the manufacture’s guidelines. Briefly, 1.8 uL diluted DNA was dried into a DNA pellet using an incubator. After that, 2.4 ul of KASP reaction mix was added (1.2 uL ultrapure water, 1.2 uL PACE® mix, 0.03 uL primers mix). The PCR programme used is a touchdown PCR and the termocycling steps conditions were as followed: pre-incubation at 94°C for 15 min followed by 10 cycles of 94°C for 20 s and then 65-75°C for 1, an additional 45 cycles of 94°C for 20 s, 57°C for 1 min. Fluorescent endpoint reading were performed on a Tecan Safire plate reader and the data analysis performed by Klustercaller software (version 2.22.0.5, LGC).

## Data Availability

The raw sequencing data has been deposited at NCBI under the accession number PRJEB83226.

## Acknowledgments

We thank Dan MacLean, Christian Schudoma, and Ram Krishna Shrestha for bioinformatics support and to Matthew Moscou for many fruitful discussions. This research was supported by the International Institute of Tropical Agriculture (IITA) and the 2Blades Foundation, and Lukas Brader Scholarship awarded to H.M.M. We also thank Norwich Bioscience Institutes (NBI) Research Computing for providing bioinformatics infrastructure support.

## Author Contributions

EGCF, YI, HMM, TN, JC, RG, GH, YY, MCA, SHB, HPvE and YKG performed research. EGCF, YI, HMM, TN, JC, RG, and YKG analysed the data. EGCF, HPvE, and YKG edited the manuscript. EGCF, and YKG wrote the paper. GM, MHAJJ, SHB, HPvE and YKG directed aspects of the project.

## Supplementary Figures

**Supplementary Figure 1.**
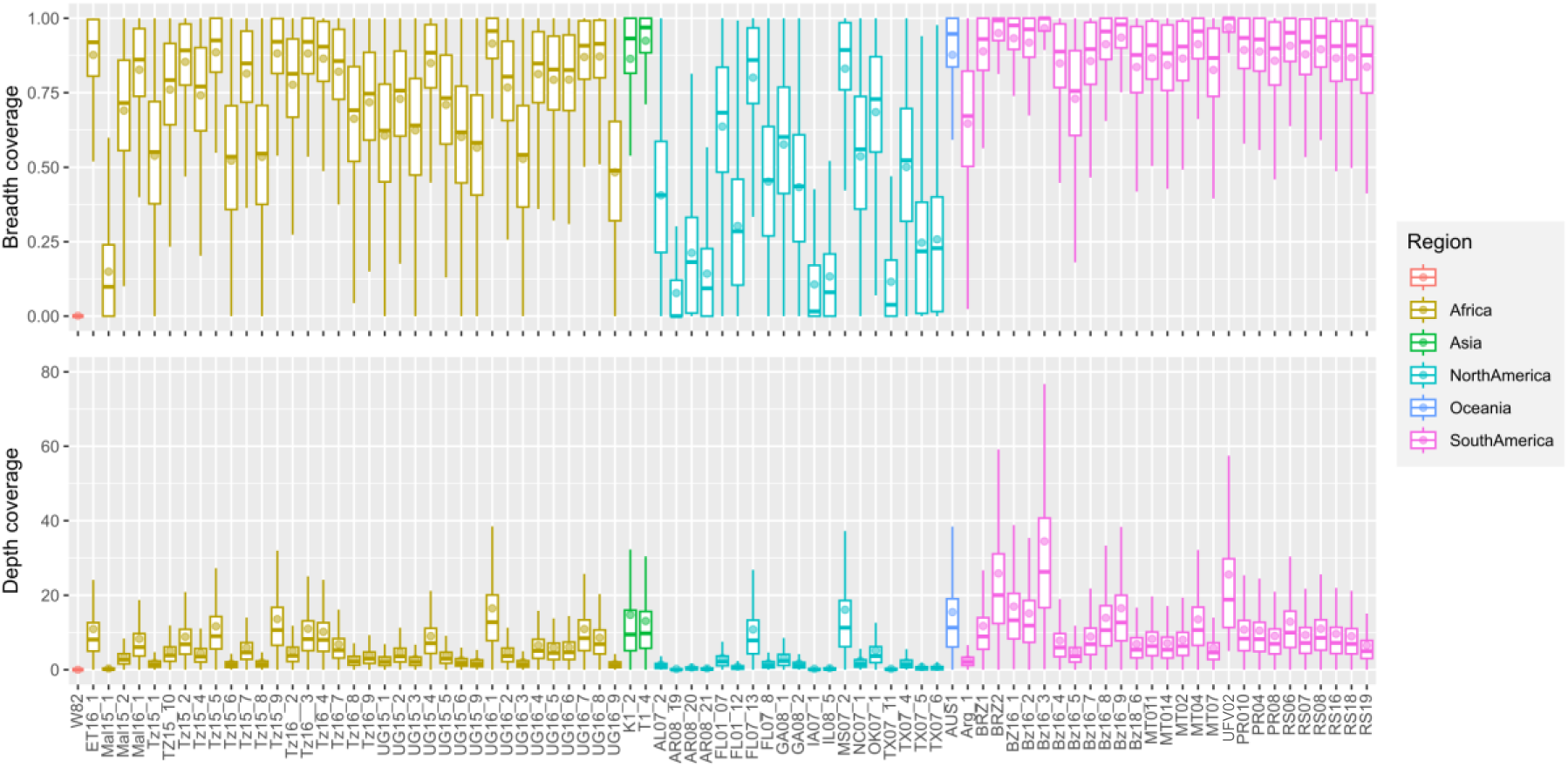
Read breadth and depth coverage of the UFV02 “Gene Catalogue” genes in 85 exome-capture samples. Read breadth and depth coverage over genic regions of 10942 protein-coding genes in the nuclear genome (exome-captured UFV02 “Gene Catalogue” genes) were calculated and represented as boxplots. Outlier points were omitted. Dots indicate mean values.

**Supplementary Figure 2.**
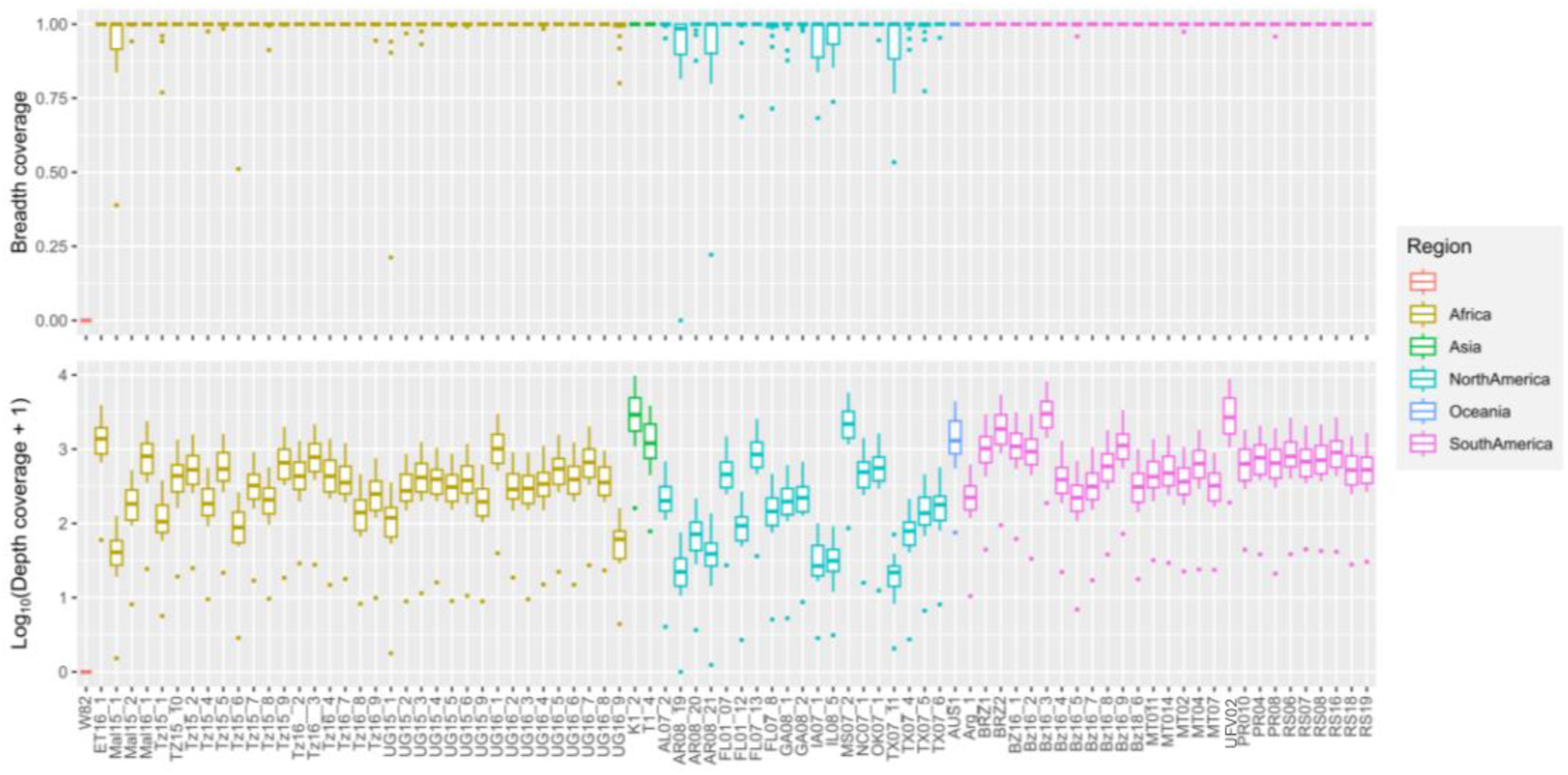
Read breadth coverage of mitochondrial genes in 85 exome-capture samples. Read breadth coverage over genic regions of 15 protein-coding genes in the mitochondria genome were calculated and are represented as boxplots.

**Supplementary Figure 3.**
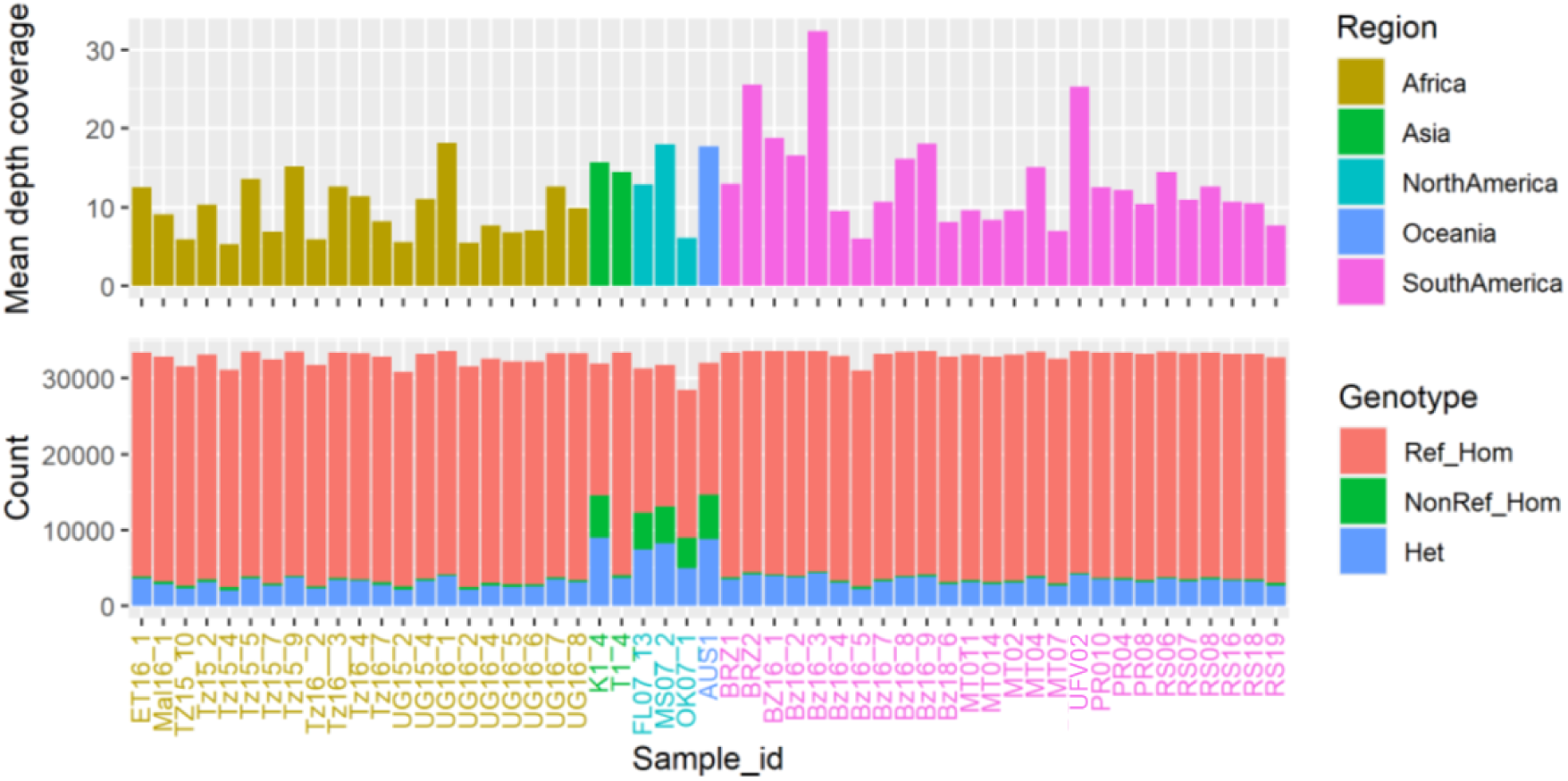
Read breadth coverage and genotypes of SNPs sites in the nuclear genome in 53 exome-capture samples.

## Supplementary Tables

**Table S1.** Information regarding the samples used for the exome-capture.

**Table S2.** Table S2 Synonymous mutations converted to the SNP markers sequences and KASP assay sequences.

**Table S3.** Genotyping results from the KASP assay COB2_2 in the new set of samples from USA and Japan.

